# Analysis of the conformational dynamics of amylose oligomers using molecular dynamics simulations

**DOI:** 10.64898/2026.07.28.741389

**Authors:** Mitsugu Araki, Biao Ma, Yukari Sagae, Katsuyoshi Masuda, Yasushi Okuno

## Abstract

Amylose contributes to starch crystallinity, but the stability of packed amylose double helices in water at elevated temperature remains insufficiently characterized. Here, we used molecular dynamics simulations to test whether chain length affects the short-timescale stability of A-type amylose oligomers in water. Six systems differing in chain length (6, 12, or 24 glucose units per chain) and oligomer size (isolated double strand or dodecamer of six double strands) were simulated, and five independent 1-μs production runs were analyzed for each simulated condition. Oligomers with six glucose units showed structural collapse accompanied by increased water penetration. By contrast, dodecamers with 12 or 24 glucose units largely retained packed double-helical organization over the simulated timescale, although fraying was observed at their ends. These results indicate that chain length and lateral packing strongly affect the early structural response of amylose-like crystalline segments in hot water. The present simulations do not establish the ultimate fate of longer oligomers at longer timescales, but they identify a relative stability difference that is relevant to molecular interpretations of hydration-driven disordering in starch.

## Introduction

Starches in rice consist of amylose and amylopectin, with their composition ratio varying depending on the type of rice. For instance, indica rice contains approximately 25% amylose and 75% amylopectin, respectively, while Japanese rice contains approximately 20% amylose and 80% amylopectin, respectively. This difference in composition ratio is thought to create variations in rice texture [1]. Amylose is a linear polymer consisting of hundreds to tens of thousands of glucose units connected together with α-1,4 glycosidic linkages. Crystallographic analysis has revealed that amylose forms single and double helical structures [2]. Furthermore, oligomeric structures with packed double helices have been reported, with packing patterns differing depending on the starch type (e.g. amyloses contained in cereals and tubers are thought to form A- and B-type packing patterns, respectively [3]). Amylopectin, on the other hand, forms a tuft-like structure through the branching of amylose molecules via α-1,6 glycosidic linkages. It contains regions with many branching points (amorphous lamellae) and regions with few branching points (crystalline lamellae) [3–5]. In the crystalline lamellae of amylopectin, the regions without 1,6-linked branches (“amylose regions”) are thought to form a three-dimensional structure similar to the crystalline structure of amylose oligomers [3].

The structural dynamics of amylose have been extensively studied using solution NMR [6] and molecular dynamics (MD) simulations [7,8,9,10,11]. Solution NMR analysis has suggested that a single-stranded amylose undergoes a transition between extended helix and compact coil structures [6]. This conformational transition has been reproduced in MD simulations [7,8], validating the force field parameters for sugar molecules. Additionally, MD simulations of a double-stranded amylose suggest that the double helical structure observed in crystallographic analysis is stably maintained in water [8]. However, the behavior of oligomeric structures with packed double helices in water is not well understood. In particular, it remains unclear how chain length and lateral packing affect the persistence of amylose-like crystalline segments in water under heating conditions relevant to starch processing.

Because branch-free segments in amylopectin crystalline lamellae are generally considered to contain approximately 12-16 glucose units [21], comparing shorter and longer oligomers may help interpret which local motifs are more susceptible to hydration-driven disordering. This study aimed to elucidate the structural dynamics of the “amylose regions” in grain starches. We specifically asked whether packed double helices with chain lengths comparable to these amylose regions remain more ordered than shorter oligomers on the microsecond timescale. Based on the oligomeric structure obtained by crystallographic analysis, we modeled double-stranded amyloses with different chain lengths, as well as oligomers with aligned double-stranded amyloses, and examined their conformational behaviors in water using MD simulations at elevated temperature. Rather than treating the simulations as a direct model of whole-granule gelatinization, we interpret them as a comparison of the relative stability of simplified local crystalline motifs in water.

## Methods

### Structure modeling of amylose

Based on the A-type crystal structure of amylose [12], we prepared six different molecular systems [(1)-(6)] that vary in chain length and the number of molecules. Water molecules were placed around the amylose molecules (Fig. 1). The type of the simulation box for molecular systems (1)-(4) was set to a truncated octahedron, while that for molecular systems (5)-(6) was set to triclinic due to their elongated shapes. The force field parameters for sugar molecules and water were set to GLYCAM06

**Figure 1.**
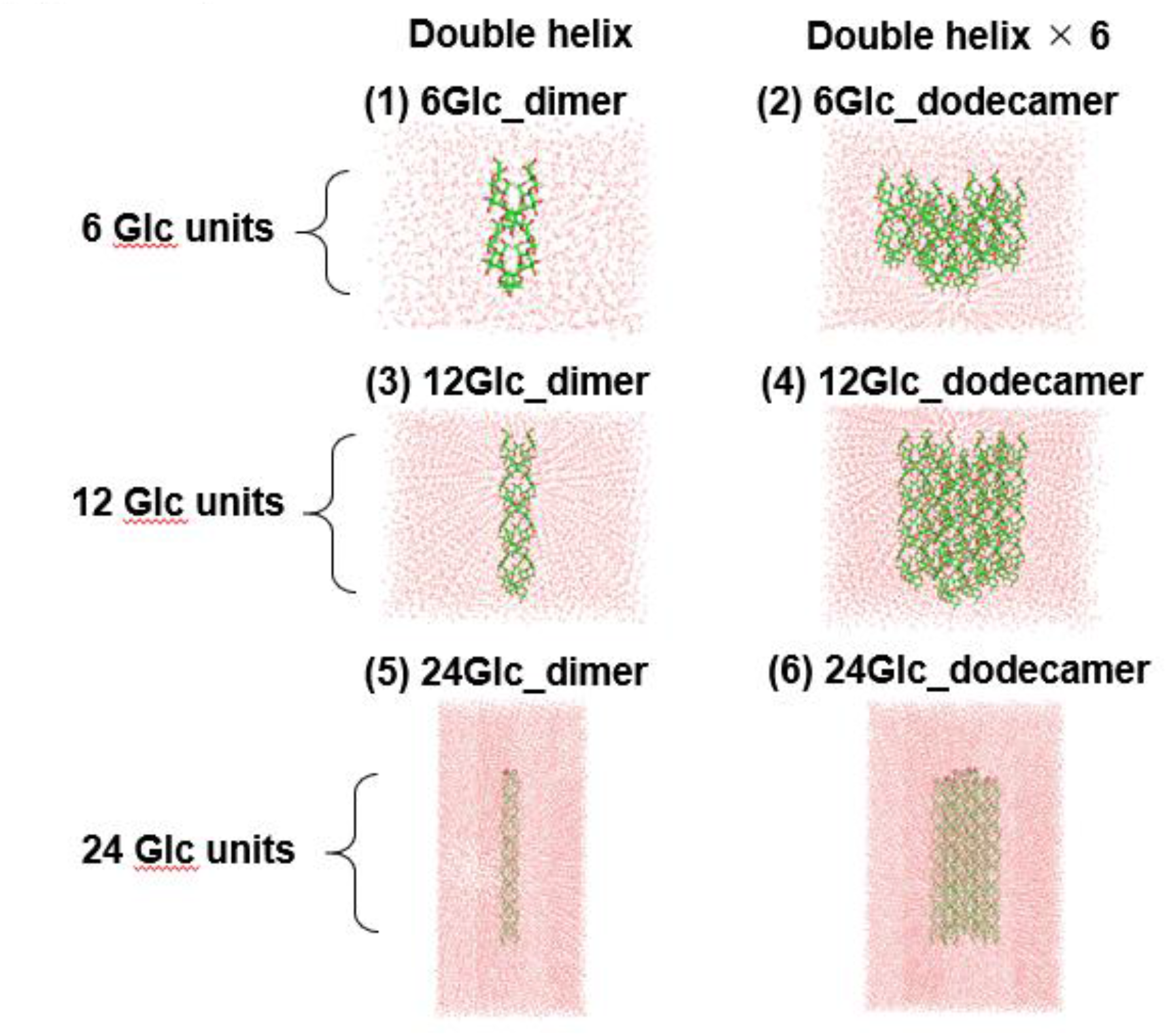
Molecular systems of amylose. Six amylose systems with different chain lengths and numbers of molecules were prepared. Water molecules were added around the amylose molecules: 2,176 (6Glc_dimer), 5,645 (6Glc_dodecamer), 6,981 (12Glc_dimer), 10,249 (12Glc_dodecamer), 24,014 (24Glc_dimer), and 38,077 (24Glc_dodecamer).

[13] and TIP3P [14], respectively. Assignment of the GLYCAM06 force field parameters to sugar molecules was performed using the doGlycans tool [15]. These systems were designed as simplified local A-type crystalline motifs and do not include α-1,6 branching points or higher-order granule architecture.

1. 6Glc_dimer, 6 glucose × 2 (one double strand)
2. 6Glc_dodecamer, 6 glucose × 12 (six double strands)
3. 12Glc_dimer, 12 glucose × 2 (one double strand)
4. 12Glc_dodecamer, 12 glucose × 12 (six double strands)
5. 24Glc_dimer, 24 glucose × 2 (one double strand)
6. 24Glc_dodecamer, 24 glucose × 12 (six double strands)

### Molecular dynamics simulations

MD simulations were performed using GROMACS 2021 [16]. Electrostatic interactions were calculated using the particle mesh Ewald (PME) method [17] with a cutoff radius of 10 Å, and van der Waals interactions were cut off at 10 Å. The P-LINCS algorithm was employed to constrain all bond lengths at their equilibrium values [18]. After each amylose system was energy-minimized, it was equilibrated for 100 ps in a constant number of molecules, volume, and temperature (NVT) ensemble and then for 100 ps in a constant number of molecules, pressure, and temperature (NPT) ensemble, with the amylose atoms held in fixed positions. Production runs were performed under NPT conditions without positional restraints. Five independent 1-μs production runs were performed with different initial velocities for each simulated condition. All analyses shown in Figs. 2–5 are based on simulations performed at 100 °C (373 K). Additional simulations at 40 °C (313 K), discussed qualitatively in the text, were used for comparison of temperature dependence and are not shown. In this series of simulations, temperature and pressure were controlled by stochastic velocity rescaling [19] and a Parrinello-Rahman barostat [20], respectively, where the time constants for temperature and pressure control were 0.1 and 2 ps, respectively. A time step of 2 fs was used in all MD runs.

**Figure 2.**
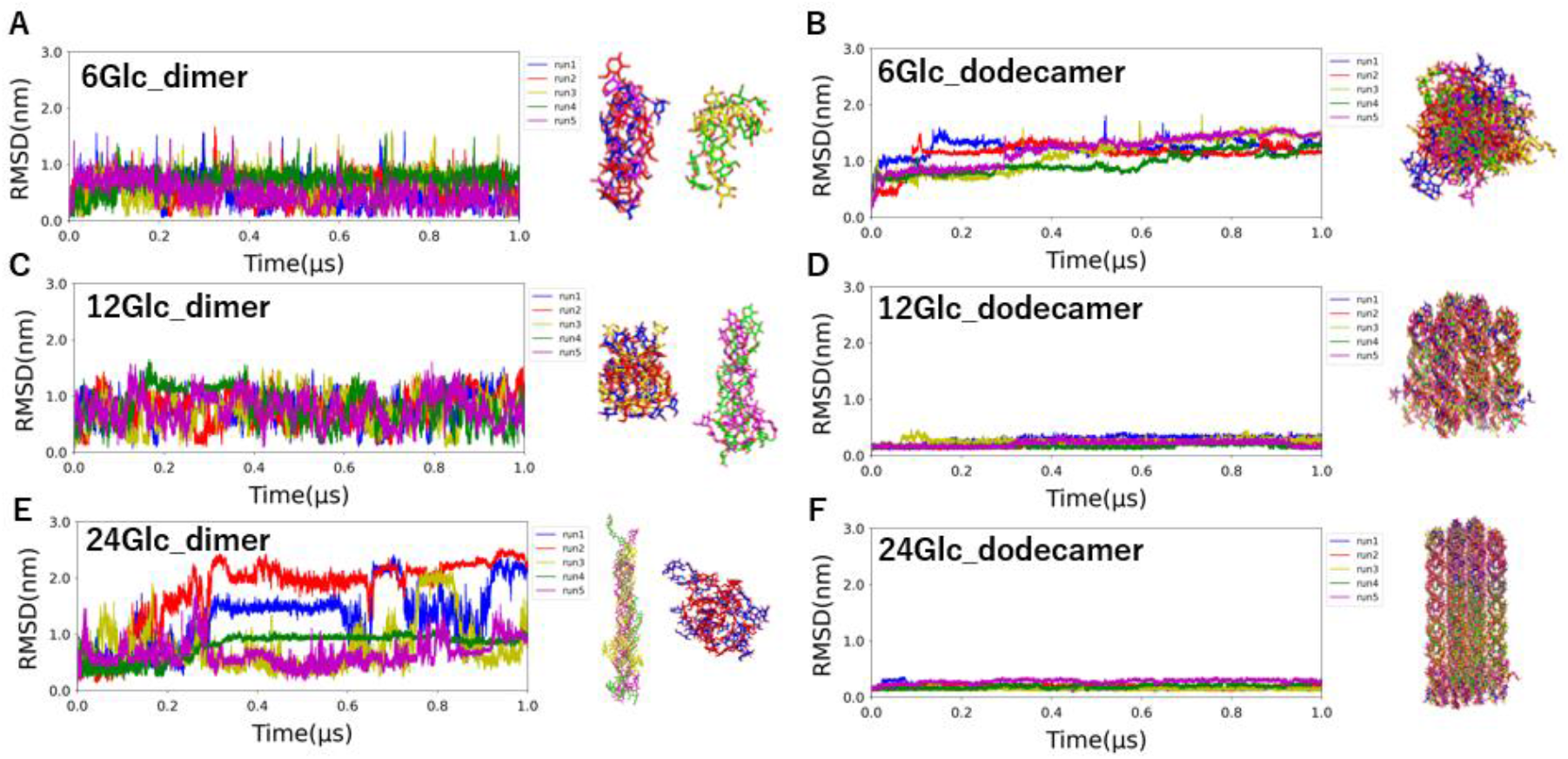
Deviation from the crystal structure at 373 K. The RMSD from the crystal structure is plotted against simulation time for five 1-μs MD simulations. The amylose structure at 1 μs is shown on the right side of each simulation, color-coded to match the corresponding graph.

## Results

### Short-chain amylose collapses more readily at 100 °C

To evaluate the behavior of amylose oligomers in boiling water, six different molecular systems with varying chain lengths and the number of molecules were prepared based on the A-type crystal structure (molecular systems (1)-(6), Fig. 1), and microsecond-scale MD simulations were conducted at 100 °C (Fig. 2). Short-chain amylose consisting of six glucose units exhibited collapse of both the double-helical structure (Fig. 2A) and a dodecamer structure with six aligned double helices (Fig. 2B). In contrast, the dodecamer structures of longer-chain amylose consisting of 12 or 24 glucose units largely maintained their overall packing over 1 μs (Fig. 2D and 2F), although fraying and local disorder were observed at the oligomer ends. The isolated 12Glc_dimer collapsed at 100 °C (Fig. 2C), and the 24Glc_dimer showed larger deviations than the corresponding dodecamer (Fig. 2E), indicating that lateral packing within the dodecamer contributes to stability beyond the intrinsic stability of a single double strand. Water molecules were observed to enter and exit the longer dodecamers through multiple side vacancies without immediate global collapse of the packed assembly (Fig. 3).

**Figure 3.**
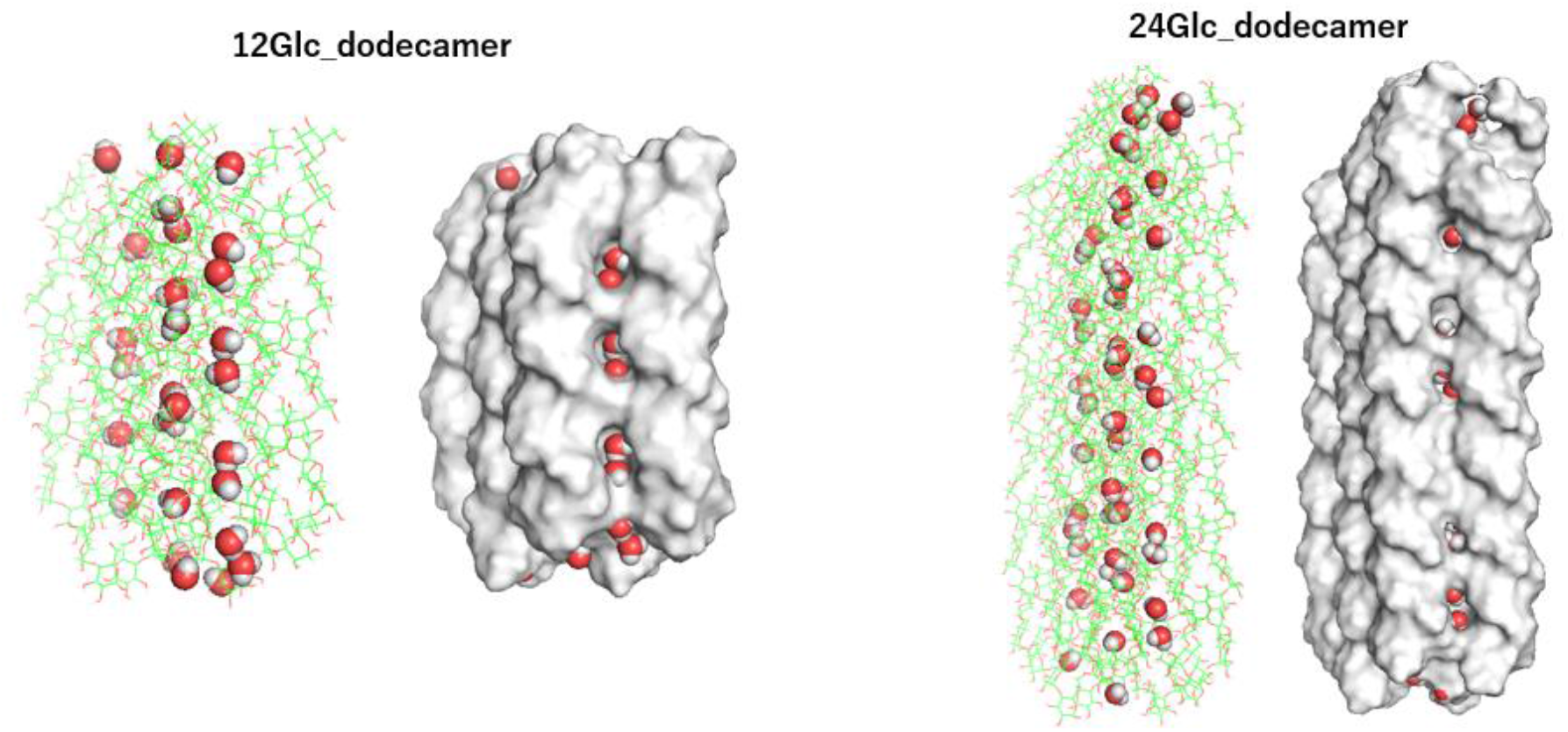
MD structures of amylose dodecamers. The final structures of the MD simulations at 373 K are presented. Amylose molecules are depicted as stick (left) and surface (right) models, with water molecules shown as spheres.

### Water penetration accompanies collapse of 6Glc_dodecamer

We further analyzed the collapse process of the dodecamer structure using the obtained MD trajectories. In a dodecamer with a short chain length (6Glc_dodecamer, molecular system (2)), individual double-helical structures collapsed concomitantly with an increase in the number of water molecules inside the amylose oligomer (Fig. 4). The temporal correspondence between increased water occupancy and the loss of helical order supports a hydration-assisted collapse mechanism in this model system. Because gelatinization has been discussed as a process in which ordered helical structures are disrupted as water penetrates starch [1], the 6Glc_dodecamer trajectory may be interpreted as a simplified molecular model for an early local disordering event rather than a complete description of macroscopic gelatinization.

**Figure 4.**
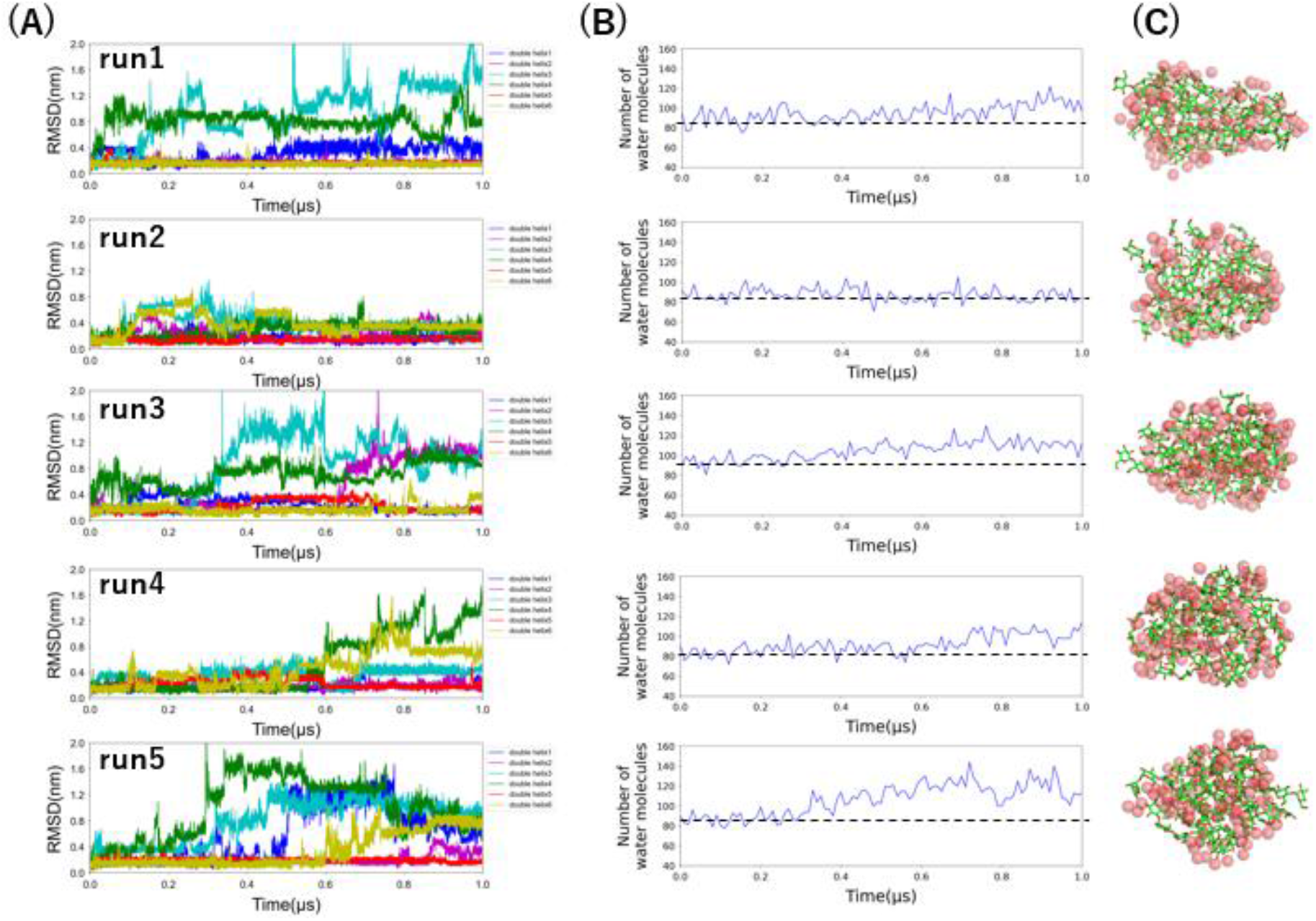
Stability of double-helical structures and distribution of water inside the amylose oligomer in MD simulations (373 K) of 6Glc_dodecamer. A) Trajectories of the main-chain RMSD. The RMSD from the crystal structure is plotted against simulation time for each of the six double strands comprising the oligomer. B) Distribution of water inside the oligomer. The number of water molecules interacting with two or more double strands is plotted. C) An oligomer structure at 1 μs. Amylose is represented as a stick model, and water molecules interacting with two or more double strands are shown as transparent spheres.

### Lateral packing stabilizes longer oligomers over the simulated timescale

In contrast, in a dodecamer with a longer chain length (12Glc_dodecamer, molecular system (4)), individual double-helical structures were largely maintained except for the ends of the oligomer, with no marked increase in the number of water molecules inside the oligomer over the simulated timescale (Fig. 5). Additional simulations at 40 °C, not shown, indicated that the isolated 12Glc_dimer remained ordered under milder conditions, whereas 6Glc_dodecamer still showed structural collapse. Together with the stability of 24Glc_dodecamer at 100 °C (Fig. 2F), these observations suggest that both chain length and lateral packing influence the rate at which ordered amylose assemblies lose their initial structure in water.

**Figure 5.**
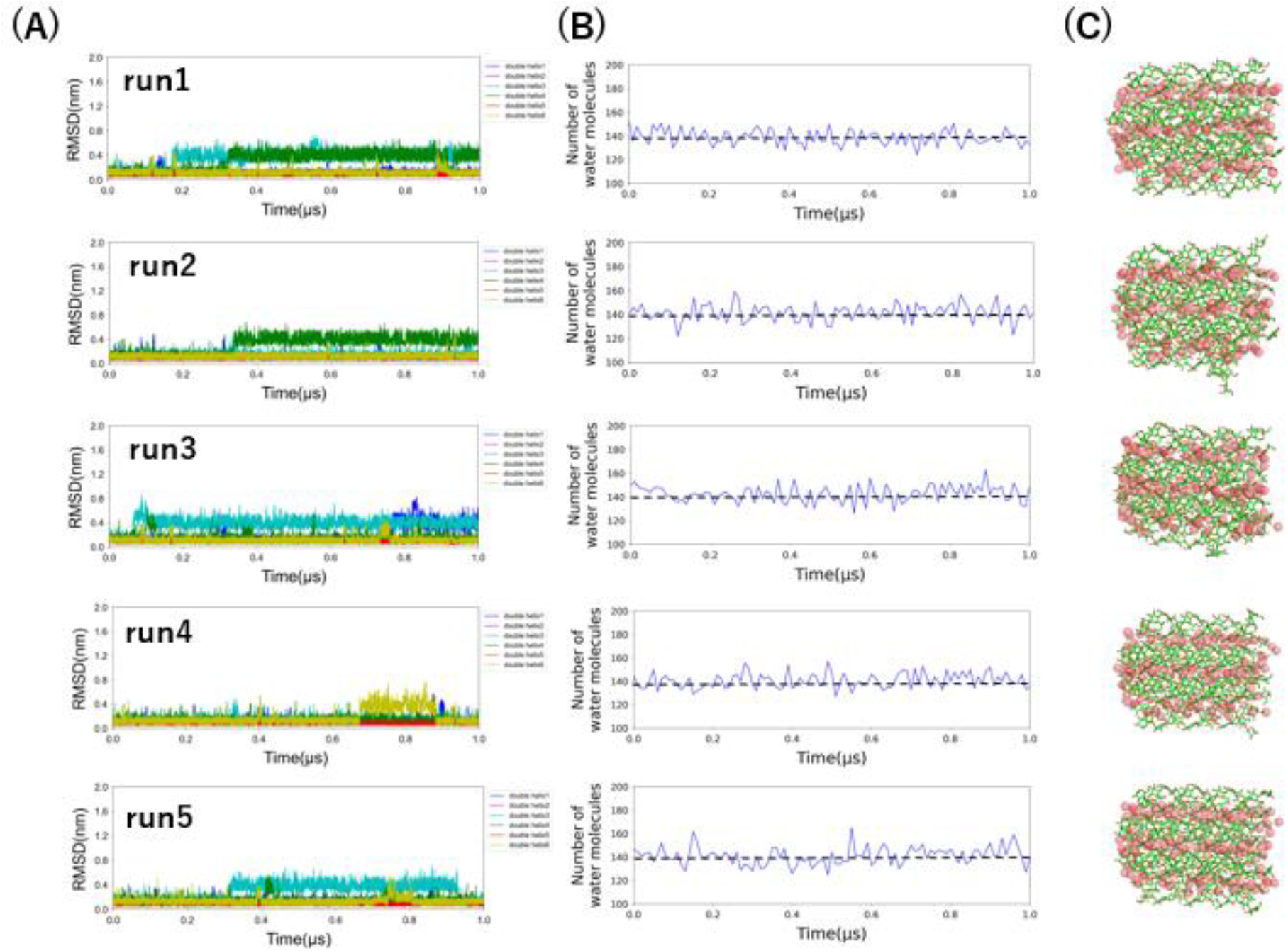
Stability of double-helical structures and distribution of water inside the amylose oligomer in MD simulations (373 K) of 12Glc_dodecamer. A) Trajectories of the main-chain RMSD. The RMSD from the crystal structure is plotted against simulation time for each of the six double strands comprising the oligomer. B) Distribution of water inside the oligomer. The number of water molecules interacting with two or more double strands is plotted. C) An oligomer structure at 1 μs. Amylose is represented as a stick model, and water molecules interacting with two or more double strands are shown as transparent spheres.

## Discussion

The main conclusion of the present simulations is therefore relative rather than absolute: longer packed oligomers are more persistent than short ones on the microsecond timescale, but we do not conclude that 12Glc_dodecamer or 24Glc_dodecamer would remain crystalline indefinitely at 100 °C. This point is important because terminal fraying is already visible in the longer oligomers. The simulations identify an order of stability under the present model conditions, not an upper limit for the time required for eventual disordering.

This interpretation provides a clearer connection to experimental starch structure. Branch-free segments in amylopectin crystalline lamellae are generally considered to have chain lengths of about 12-16 glucose units [21]. The relative persistence of the 12Glc and 24Glc dodecamers is therefore consistent with the idea that local crystalline regions in cereal starch can remain ordered longer than very short oligomers when exposed to hot water. At the same time, real starch gelatinization is a multiscale process influenced by amylopectin branching, crystal polymorphism, amorphous regions, water transport, and granule architecture [3–5]. The present models represent only isolated A-type amylose-like motifs in bulk water.

Several limitations should be noted. First, the models do not include α-1,6 branching points or interactions with neighboring amorphous regions. Second, the principal analyses are limited to 1-μs trajectories at 100 °C, and slower disordering events may occur beyond this timescale. Third, the present work does not directly predict macroscopic gelatinization temperature, viscosity change, or the behavior of full starch granules. Even with these limitations, the simulations show that water penetration and structural collapse are strongly chain-length dependent in this simplified setting, and that lateral packing can stabilize amylose double helices that are less stable when isolated.

## Conclusion

In conclusion, the present MD simulations show that A-type amylose oligomers with six glucose units are markedly more susceptible to hydration-associated structural collapse than packed oligomers with 12 or 24 glucose units over the simulated timescale. The most defensible interpretation is not that longer oligomers are permanently stable in boiling water, but that they undergo disordering more slowly than short oligomers under the present conditions. This chain-length-dependent difference provides a concrete molecular hypothesis for how local crystalline amylose-like segments respond to heating in water. Future work extending the simulations in time and incorporating branched amylopectin motifs should help test how broadly this relative stability ranking applies in more realistic starch models.

## Abbreviations

MD: molecular dynamics
RMSD: root-mean-square deviation.

## Data availability

The input files, molecular dynamics trajectories, and processed data supporting the findings of this study are available from the corresponding authors upon reasonable request. Because of file size, these data are not deposited in a public repository.

## Acknowledgements

The simulations were performed on the Fugaku supercomputer provided by the RIKEN Center for Computational Science through the HPCI System Research Project (project IDs: hp230216 and hp240211).

## Funding

This study was supported by the Ministry of Education, Culture, Sports, Science and Technology (MEXT, Japan) as “Program for Promoting Researches on the Supercomputer Fugaku” (Simulation- and AI-driven next-generation medicine and drug discovery based on ‘Fugaku’, JPMXP1020230120) (to Y.O.). M. Araki and K. Masuda were supported by Zensho Holdings Co., Ltd.

## Author contributions

K.M. and Y.O. conceived and supervised the study. M.A. and B.M. performed the molecular dynamics simulations. M.A. and Y.S. analyzed the simulation data. M.A. and K.M. wrote the manuscript. All authors discussed the results, edited the manuscript, and approved the final version.

## Additional information

Competing interests: The authors declare no competing interests. Correspondence and requests for materials should be addressed to K.M. or Y.O.

## Notes

### Competing Interest Statement

The authors have declared no competing interest.

